# Mechanical Dissociation of Tissues for Single Cell Analysis Using a Simple Motorized Device

**DOI:** 10.1101/2023.05.03.539271

**Authors:** Mayowa Amosu, Andrew J. Gregory, John D. Murtagh, Nitay Pavin, Carson Taylor Meyers, Juan Grano de Oro Fernandez, Kaitlyn Moore, Katharina Maisel

**Affiliations:** Fischell Department of Bioengineering, University of Maryland, College Park, MD 20742; UMD Terrapin Works, University of Maryland, College Park, MD 20742

**Author notes:** Correspondence to: Dr. Katharina Maisel, Fischell Department of Bioengineering, University of Maryland, College Park, MD 20742, United States. **Contributions:** MA, AG, and JM collaborated on mechanical engineering, prototyping, and device design. NP and CT wrote and debugged all coded device software. MA led all major experiments and data analysis; JG and K. Moore assisted in experiments and experimental analysis. K. Maisel supervised experimental planning, data analysis and manuscript drafting. MA, JG, K. Moore and K. Maisel drafted, edited, and finalized the manuscript.

## Abstract

The use of single cell analysis methods has grown rapidly in the last two decades and has led to rapid discoveries in cell biology and beyond. Single cell analysis requires complex systems like tissues to be dissociated, separating individual cells from extracellular tissue materials. This requires manual processing of tissues and materials through chopping, pipetting, and suspension with enzymes for degradation of the structural elements of the tissue. Manual processing can be time consuming and lead to variability between scientists. Automating this process through motorized dissociation could thus improve reproducibility of research and reduce time of cell manipulation prior to analysis. Here, we have designed a low-cost, customizable automatic tissue dissociator device that can be easily assembled by research groups for individual use. Our device allows for customizable programmed dissociation protocols for ease of use and reproducibility between researchers and can be placed into heat or cold environments based on the protocol need. We have found this device comparable in cell viability and reproducibility to manual dissociation, while significantly reducing time spent and even enhancing cells extracted from more fibrous tissues. Broad dissemination and use of this device could enhance single cell analysis reproducibility and provide a time-saving alternative to the currently used manual dissociation protocols.

## INTRODUCTION

Single cell analysis has rapidly become crucial for new biomedical discoveries, whether this is for applications like flow cytometry, used to identify different cell types, or single cell sequencing, used to identify genomic or transcriptomic variation between cells (1). To perform such cell isolations from tissues of interest requires mincing dissected tissues and pushing them through a fine cell strainer to filter out connective tissue from the desired cells (**Figure 1A**). Isolating adherent cell types, like dendritic cells or macrophages, or cells from particularly fibrous tissues, requires additional mechanical or enzymatic separation steps (2–4). This process is generally done manually, making it highly time consuming and prone to user variability for assessment of cell yields and sample viability. Therefore, it is crucial to introduce customizable options for automated tissue dissociation. While some attempts have been made to design such systems, the existing options are not always readily accessible, particularly in academic labs and lower resource settings, largely due to the cost-prohibitive nature of these devices (5). Furthermore, these devices are not always customizable to individual needs of a research group (6). Here, we have designed a tissue dissociator device aimed to automate digestion of whole tissues or tissue pieces into single cell suspensions with aid of digestive enzymes and mechanical disruption. This device can be easily assembled in the lab, placed into heating or cooling chambers for temperature regulation, customized for the required number of tissues to dissociate, and programmed with desired dissociation protocols. Broad use of this device could significantly improve reproducibility in cell extraction protocols and provide a time saving alternative to manual dissociation.

**Figure 1.**
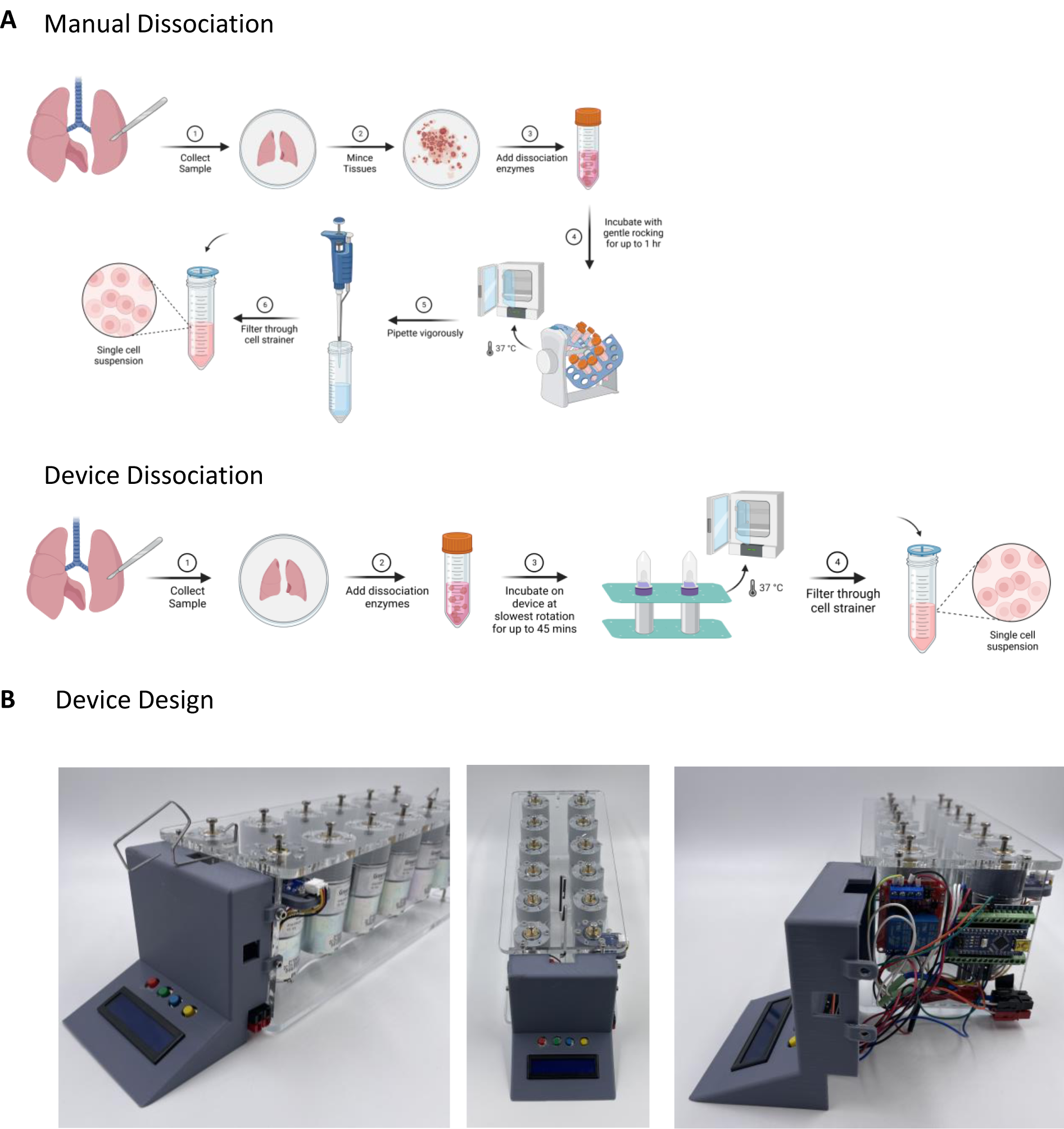
Process Diagrams. A) Steps in the process of preparing single cell suspension using simple dissociator device vs manual dissociation. B) Dissociator device design and features.

## DEVICE SUMMARY

Our main design allows for simultaneous digestion of up to 12 tissues through an automated process. The device is composed of 12 individual motors wired in parallel and powered by a standard wall plug through an AC/DC adapter with an adjustable voltage dial to control the rotation/speed of the motors. The motors are turning a hex bolt that fits snuggly into the top of Miltenyi Biotec’s C-tubes (Miltenyi Biotec, 130-093-237). The C-tubes are held in place by downward tension on an acrylic plate that latches on either side to the top plate where the motors are secured (**Figure 1B**). Because the motors are wired in parallel, their speed at any given voltage should not vary much, but the load (number of C-tubes mounted on the device) will affect speed even when voltage is kept constant. To measure rotations per minute (rpm), we have incorporated a tachometer using a hall effect sensor and a fixed magnet to one of the motor shafts **(Figure 2C)**. To reverse the direction of rotation, we have also included a programmable switch that reverses the positive/negative charges to the motors All these features are integrated using coded software (Arduino IDE Software) on an Arduino Nano. Using connected buttons and an LCD panel **(Figure 3A)**, it is possible to create and run saved and custom protocols, automatically reverse the rotational direction at specified times of a protocol, and display the current motor speed and time left to complete a programmed protocol (**Figure 5**).

**Figure 2.**
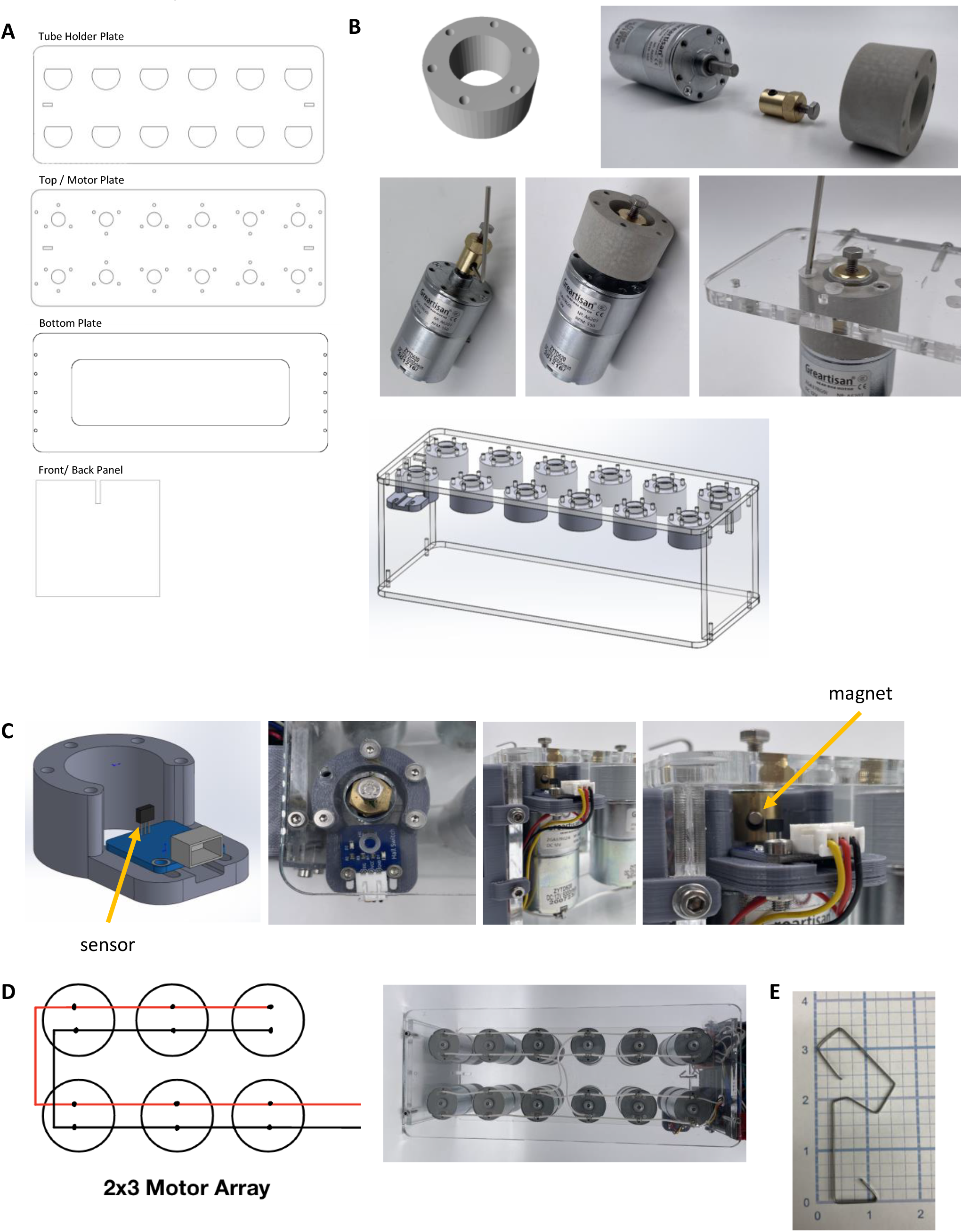
Motor Assembly. A) Images of acrylic sheet CAD files B) Assembly of motor/ coupler/ hex bolt / motor spacer unit onto the motor plate C) RPM Sensor D) Motor wiring

**Figure 3.**
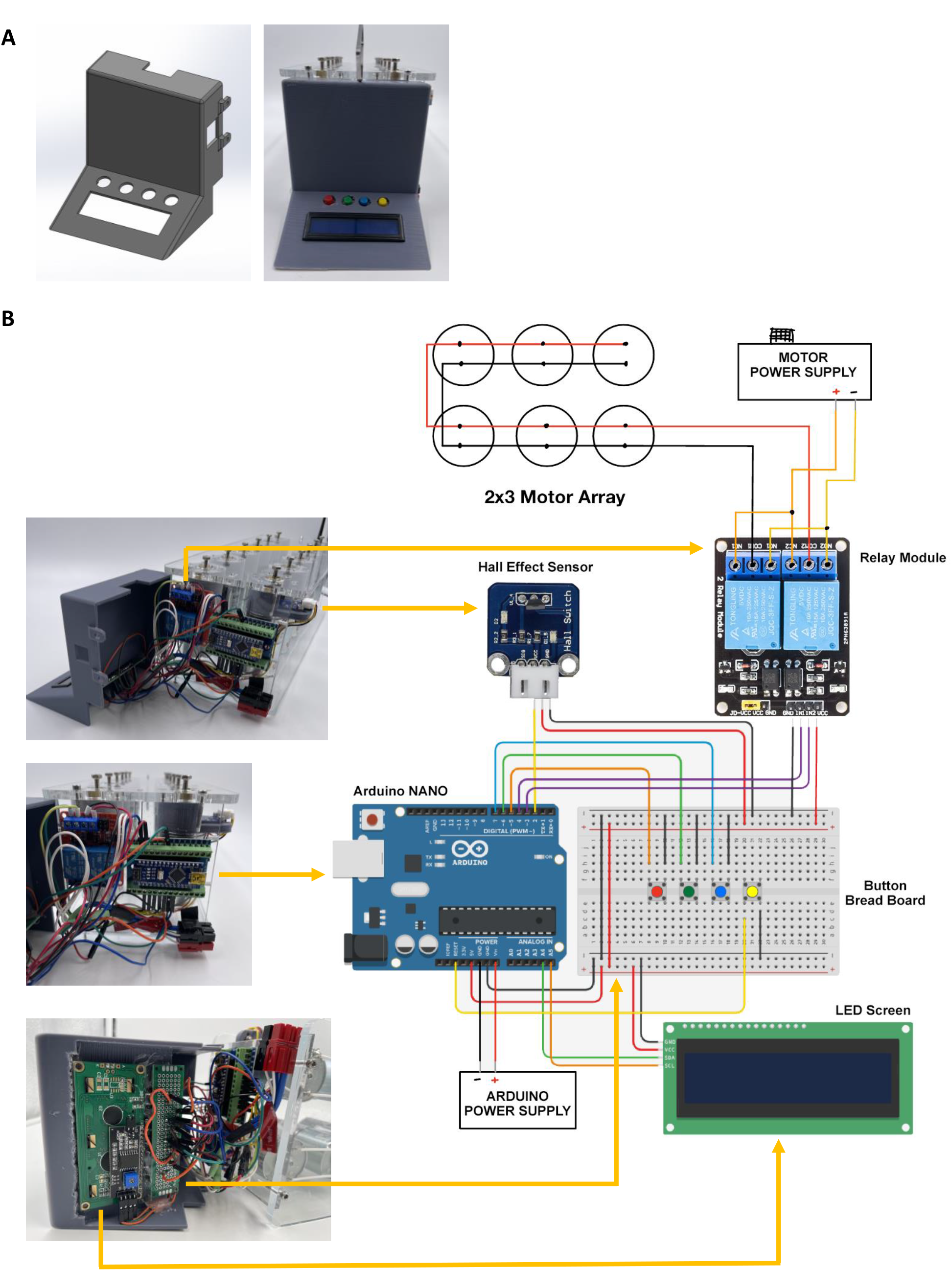
Control Panel Assembly. A) Image of Control Panel CAD file B) Control panel / Power Supply wiring diagram

## METHOD DETAILS

### MATERIALS

See Table 1 for detailed list of assembly and operation materials.

**Table 1:**
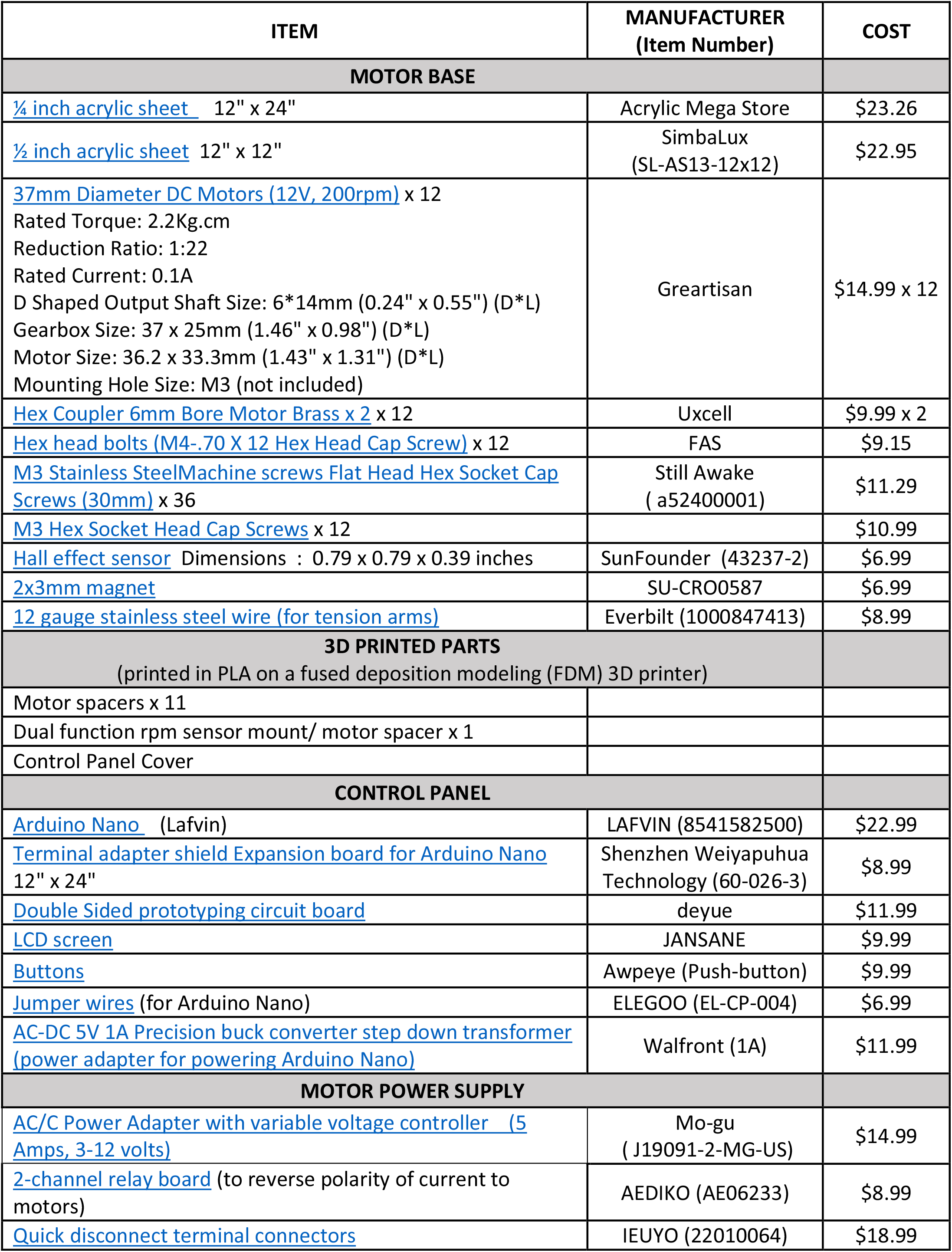
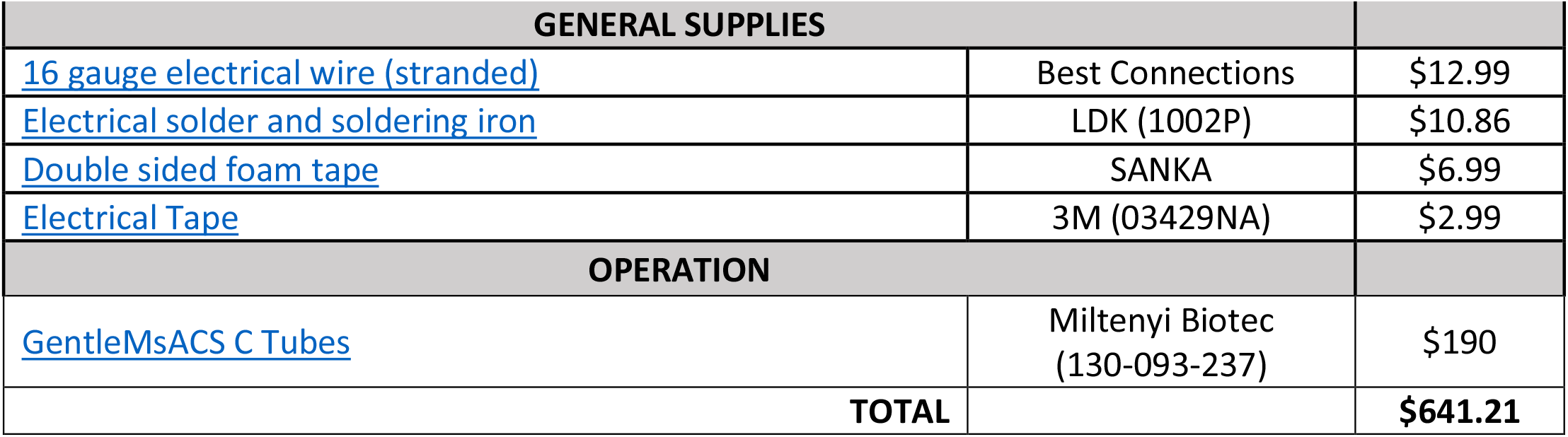
Required materials for assembly and operation of device components.

### DEVICE FABRICATION AND ASSEMBLY: MOTOR ARRAY

The device container is made of acrylic sheets (¼ and ½ inch) laser cut to fit 12 motors (**Figure 2A**, CAD files for configurations of 2, 4, 6, 8, 10, and 12 motors are available in the Supplemental Materials). The original design of the top sheet has spaces for 12 motors in 2 rows of 6, but the number of motors can be customized to fit desired needs and incubator space requirements. Motor spacers (**Figure 2B**) are 3D-printed from polylactic acid (PLA) filament on a fused deposition modeling (FDM) 3D printer and are necessary to mount the motors at the correct distance from the acrylic top sheet that accommodates the 6mm motor coupler and an M4 hex head bolt (**Figure 2B**). One motor spacer has dual function as a mount for a hall effect sensor used to detect motor revolutions for rpm calculations (**Figure 2C**).

To fabricate the structural components of the motor and frame, follow the instructions below:

1. Laser cut the top and bottom device panels from ¼ inch acrylic sheets using the CAD design of your choosing (2-12 motors)
2. Laser cut the front and back device panels from ½ inch acrylic sheets
3. Print motor spacers (1-11) and hall effect sensor mount spacer (**Figure 2C**) from PLA on an FDM 3D printer.
4. Connect the 6mm brass coupler to the D-shaped shaft of the motor and secure it in place with the provided M4 set screws. (**Figure 2B**).
5. Add Loctite into the top of the brass coupler before screwing in the M4 hex bolt. (The Loctite prevents the bolt from coming undone).
6. Place a motor spacer over the hex bolt and insert M3×30mm flat head machine screws in the holes provided on the top sheet to secure it directly to the motor below.
7. Repeat steps 1-3 for (n-1) number of motors as required for your device needs. *Note: the positive/negative motor electrical connections should be installed in the same orientation such that all the positive connections are on the same side of the device. This facilitates easy wiring later (****Figure 2D***).
8. For the dual-purpose motor spacer, place a 2×3mm magnet over the metal set screw in the side of the brass coupler and secure the hall effect sensor along the track. The sensor should be close enough to the magnet to trigger the switch (indicated by a lit bulb), but far enough away not to interrupt the motor rotation or dislodge the magnet while in motion.
9. Once the magnet and sensor are in place, repeat step 3 for the dual-purpose motor.
10. Once all motors are assembled on the top sheet, flip device over and wire polarity of motors in parallel according to diagram (**Figure 2D**).
11. Back, front, and bottom panels can now be assembled:

a. Drill holes for 2 M3 × 12mm screws into the top and bottom of both ½ acrylic back and front panels.
b. Drill holes for 2 M3 × 6mm screws on either side of the front panel. (These will be used to secure the control panel later).
c. Attach the top plate with the wired motors to both front and back panels using 4 M3 × 12mm screws.
d. Attach the bottom plate to both front and back panels using 4 M3 × 12mm screws.
12. Bend 3mm flat wire using pliers into shape to make the tension arms (**Figure 2E)** and insert into the slots in the front/back panels and hook into the slots provided at either end of the top motor plate (**Figure 1B/2A**).

### DEVICE FABRICATION AND ASSEMBLY: CONTROL PANEL & POWER SUPPLY

The device control panel is 3D-printed PLA casing that houses the internal electronic components. The control panel creates an interface between the software programmed into the Arduino Nano and the hardware (LCD screen, button, motors, and hall effect sensor). Instructions on the printing and assembly of the control panel and power supply can be found below:

1. Print the control panel (**Figure 3A**) casing in PLA on an FDM 3D printer.
2. Wire the electrical components using jumper wires and electrical wire according to the wiring diagram (**Figure 3B**) in this suggested order:
  a. Screen (connected to the Arduino Nano and grounded on the button bread board)
  b. Buttons (fitted to bread board and connected to Arduino Nano)
  c. Relay module (connected to the Arduino Nano, the motors, and the variable voltage power supply)
  d. Hall effect sensor (connected to the Arduino Nano and grounded on the button board)
  e. Arduino Nano (connected to all other components and its own power supply) *Note: The diagram shows two separate power supplies for the motors (variable voltage) and Arduino Nano because it is not recommended to subject the Arduino Nano to variable voltage. The quick disconnect terminal adapters are an optional feature if the user would prefer easy disconnect of the device from the power sources for ease of transport*.
3. Mount the screen and colored buttons into the cutouts on the control panel casing and secure using double-sided foam tape.
4. Load software code onto the Arduino Nano:
  a. Install and open the program “Arduino IDE” on a computer (https://www.arduino.cc/en/software).
  b. Connect Arduino Nano into the computer via USB.
  c. Select Arduino Nano from the drop-down menu at the top of the screen.
  d. Open the file “DeviceCode_2023.ino” containing the software code in Arduino IDE.
  e. Load the code onto the connected Arduino Nano by clicking the “Upload” button.
5. Mount the Arduino Nano to the expansion board and secure to the front panel using double-sided foam tape. Ensure that the position of the USB Mini-B port is in line with the cutout on the right side of the control panel (**Figure 3A**). This allows users to upload modifications to the software code after assembly is complete.
6. Secure the relay module to the front panel using double-sided foam tape.
7. Ensure that wire connections are tight and secure before placing the control panel cover over the electrical components and securing in place with 2 M3×6mm screws on either side of the front panel.

## DEVICE OPERATION

### Loading Samples

1. Connect the Arduino Nano to power supply (the LCD screen will light up automatically).
2. Connect the motors to power supply and ensure that the voltage control on the power adapter is turned off (the motors should not be turning).
3. Prepare tissue samples by chopping material to pieces small enough to fit under the rotor in the Miltenyi Biotec gentleMACS C Tubes.
4. Ensure proper closure of tubes by rotating cap clockwise until you feel a “click”.
5. Install tubes upside down onto motor driven hex bolt heads.
6. Fit the tube holder plate (**Figure 2A**) over the D-shaped tube bottoms.
7. Secure tubes to motor plate by latching tension arms into the acrylic plate (**Figure 4A**).
8. Turn the voltage control on.
9. The device and samples are now ready to run a protocol/program (the motors will begin turning once you have selected a mode or protocol).

**Figure 4.**
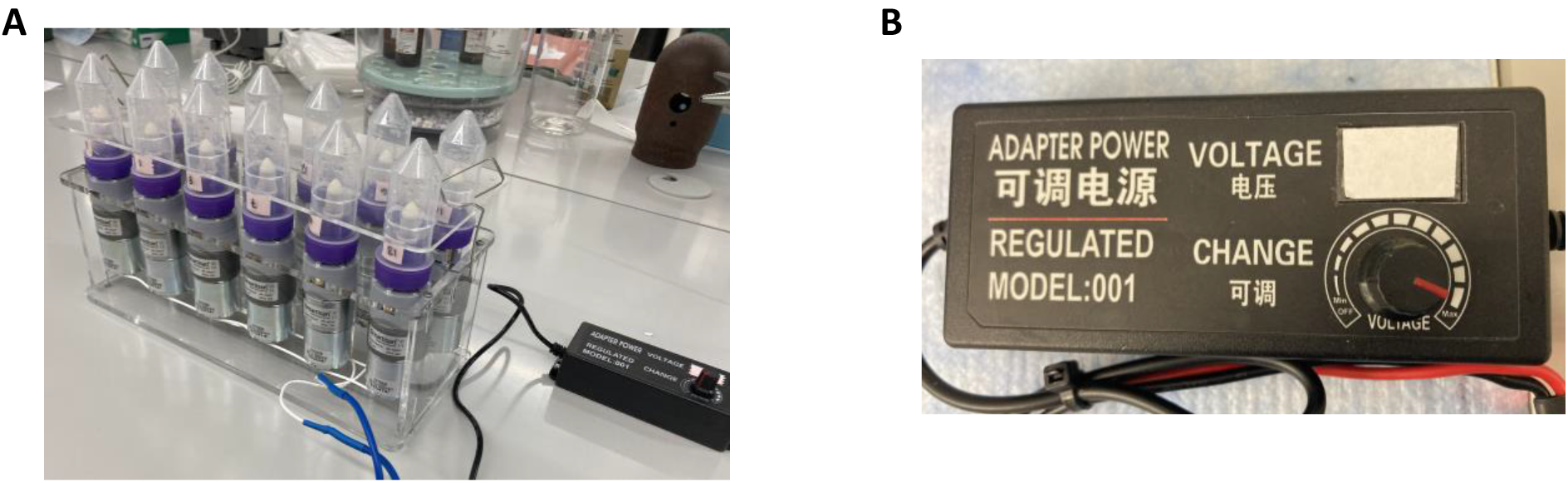
Device Operation. A) gentleMACS C Tubes loaded onto device and secured with tube holder plate and tension arms B) Variable voltage regulator with dial

**Figure 5.**
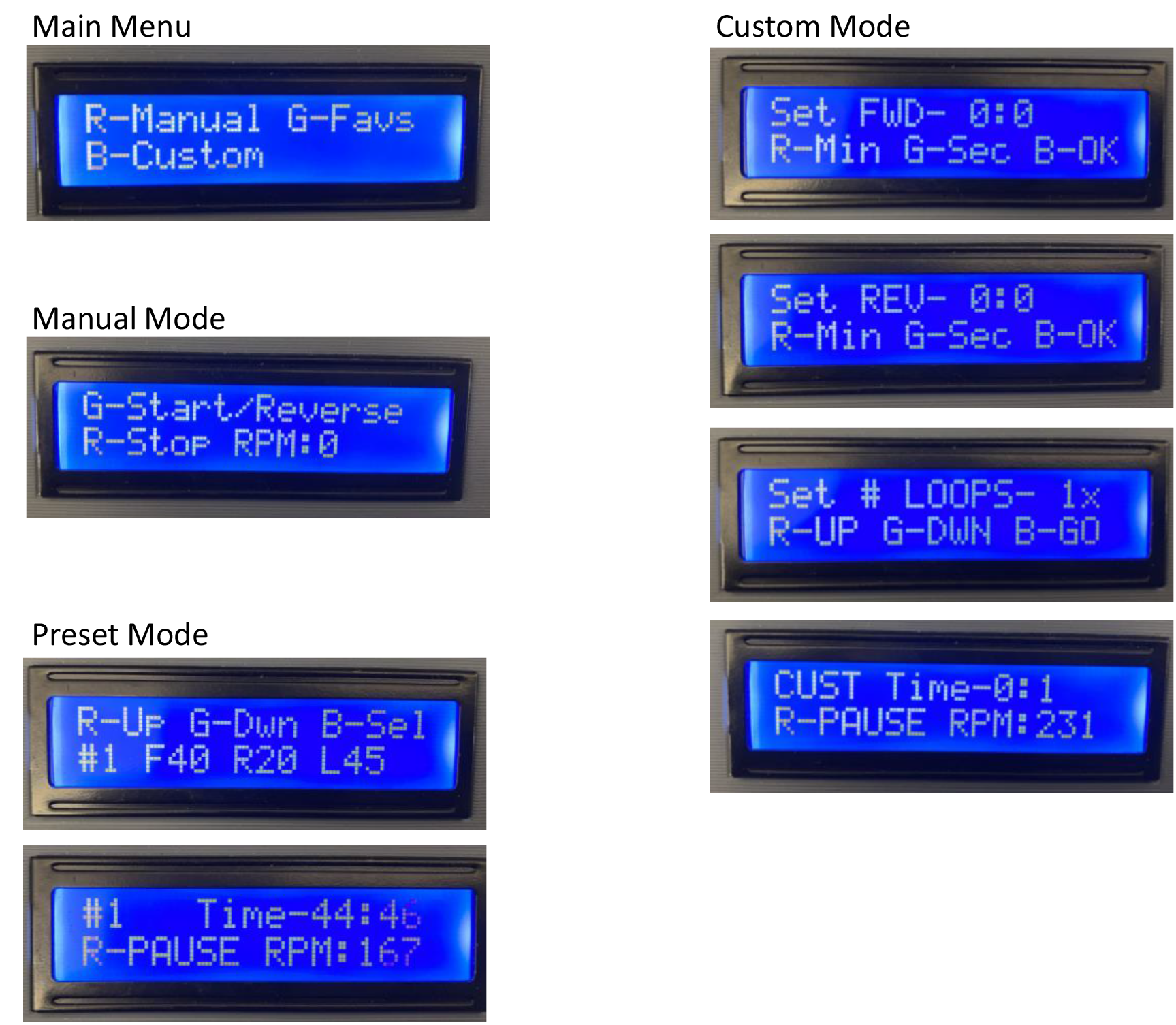
Software Operation – Control Panel Modes. A) LCD Screen Displays for various modes of device operation

### Operating Software

The main menu allows you to choose between 3 modes of operation by pushing 1 of 3 colored buttons: Red: Manual, Green: Preset Programs, Blue: Custom Programs, and Yellow: Resets the Arduino Nano and returns you to the main menu. In the software menus, the letter R, G, and B refer to the color button that corresponds to a particular selection (**Figure 4B**).

R = Red button

G = Green button

B = Blue button

Once a program is selected and the motors are turning, the LCD screen will display calculated RPM values in the bottom right corner (**Figure 5A**). The speed of the motors can be adjusted to the desired RPMs using the dial to control voltage (**Figure 4C**). When running preset and custom programs, the remaining time in the current program is displayed in the top right corner.

### In Manual Mode

- R = Pause (Stops the motors from turning)
- G = Run (Starts the motors)

### In Preset Mode

- R = Up (Cycles through preset program in ascending order #1 to #9)
- G = Down (Cycles through preset program in descending order #9 to #1)
- B = Select and Run (Begins the selected program)

Preset programs may be adjusted by entering values directly into the Arduino IDE software code and reloading the updated code to the Arduino Nano. To add your own preset programs, go to lines 54-58 in the code. There are 10 “slots” available for preset programs with 3 adjustable components:

1. Duration of forward rotation in seconds
2. Duration of reverse rotation in seconds
3. Number of times to “loop” or repeat steps 1 and 2

In the sample code for preset programs below, each position separated by a comma, corresponds to values for an individual preset program up to 10:

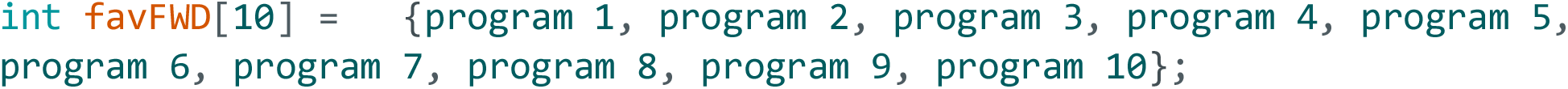

In the code below, **preset program #1 is coded as follows**:

Step 1. Forward Rotation: **30** seconds

Step 2. Reverse Rotation: **10** seconds

Step 3. Loop Steps 1-2: **4** times

And **preset program #2 is coded as follows**:

Step 1. Forward Rotation: **300** seconds

Step 2. Reverse Rotation: **20** seconds

Step 3. Loop Steps 1-2: **9** times

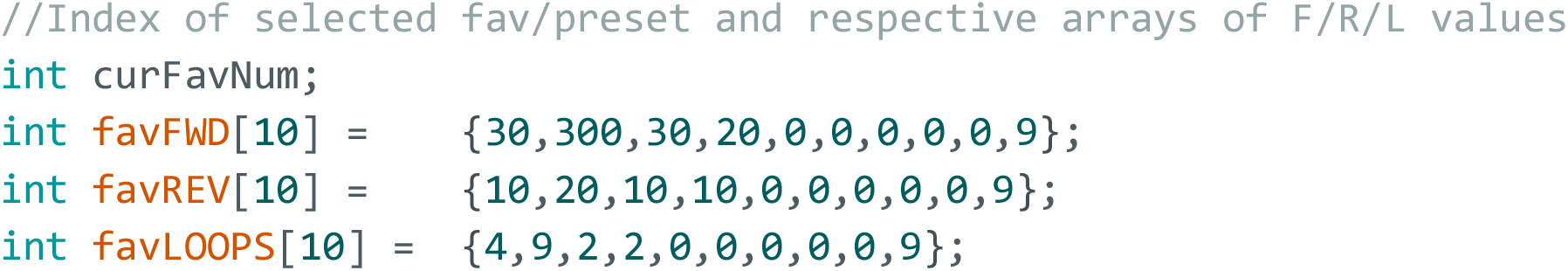

### In Custom Mode

You will move through 3 different menus to dictate to the software the specifications of the protocol you would like to run.

Menu #1: Duration of Forward Rotation

- R = Seconds
- G = Minutes
- B = Select

Menu #2: Duration of Reverse Rotation

- R = Seconds
- G = Minutes
- B = Select

Menu #3: Loop (number of times to repeat Steps 1-2)

- R = Increasing integers
- B = Select and Run (Begin the selected program)

Custom programs are not retained in the device memory but can be programmed as one of the preset programs.

## QUANTIFICATION AND STATISTICAL ANALYSIS

### Device Validation

We prepared side-by-side single cell suspensions using both mechanical-enzymatic tissue dissociation with our device and manual-enzymatic tissue dissociation to determine the differences, if any, in cells recovered for downstream applications. We compared total cell yields per tissue, percent cell viability, and used flow cytometry to compare potential differences in surface marker expression.

Data were analyzed using Graphpad Prism (GraphPad, La Jolla, CA). Unpaired Welch t tests were used to compare pairs of samples or groups with sample sizes n>4 mice representing 2 replicate experiments. Differences were considered statistically significant when p < 0.05. *p<0.05, **p<0.01, ***p<0.001, ****p<0.0001.

### Materials

- DMEM (**Dulbecco’s Modified Eagle Medium**)
- Digestions Enzymes

∘ Collagenase 4 (Worthington, CLS4 LS004188)
∘ Collagenase D (Roche, 11088866001)
∘ DNAse (Roche, 11284932001)
EDTA
Red Blood Cell Lysis Buffer
gentleMACS C Tubes (Miltenyi, 130-093-237)

### Methods

Protocol used for **manual** dissociation

1. Remove desired tissue and keep in cold media (DMEM with 5% FBS).
2. Mince tissue using sharp dissection scissors as finely as possible.
3. Transfer to 15mL conical tube with 5mL media.
4. Add digestion enzymes according to the tissue and cell types being isolated.
  a. For the presented experiments, we used Collagenase 4 (1 mg/mL), Collagenase D (1 mg/mL), and DNAse (40 μg/mL).
5. Incubate on shaker for 1 hour at 37°C.
6. Pipette sample up and down 100 times.
7. Add EDTA to bring the sample volume to 5mM concentration.
8. Pipette sample up and down 100 times.
9. Pipette sample up and down 100 times vigorously.
10. Push sample through a 70μm cell strainer.
11. Spin collected cells down using a centrifuge at 300g for 5 minutes.
12. Discard supernatant and resuspend cells in 1mL red blood cell lysis buffer.
13. Incubate for 1 minute.
14. Quench lysis buffer with 9mL cold media.
15. Spin collected cells down using a centrifuge at 300g for 5 minutes.
16. Discard supernatant and resuspend cells in buffer or media.
17. Cells are ready for downstream applications.

Protocol used for **mechanical** dissociation:

1. Remove desired tissue and keep in cold media (DMEM with 5% FBS).
2. Chop tissue using sharp dissection scissors (does not need to be finely minced).
3. Transfer to C tube and add 1mL media.
4. Load C tube on device and run program at 200rpm: Forward 30 seconds, Reverse 10 seconds, Loop 4 times.
5. Add digestion enzymes in 4mL media according to the tissue and cell types being isolated.
  a. For the presented experiments, we used Collagenase 4 (1 mg/mL), Collagenase D (1 mg/mL), and DNase (40 μg /mL).
6. Load C tube on device and run program at 200rpm: Forward 30 seconds, Reverse 10 seconds, Loop 4 times.
7. Transfer device with tubes still loaded to 37°C incubator for 45 minutes.
  a. Run program at 30rpm at 37 ° C: Forward 270 seconds, Reverse 30 seconds, Loop 9 times.
8. Add EDTA to bring the sample volume to 5mM concentration.
9. Load C tube on device and run program at 100rpm: Forward 30 seconds, Reverse 10 seconds, Loop 2 times.
10. Push sample through a 70μm cell strainer.
11. Spin collected cells down using a centrifuge at 300g for 5 minutes.
12. Discard supernatant and resuspend cells in 1mL red blood cell lysis buffer.
13. Incubate for 1 minute.
14. Quench lysis buffer with 9mL cold PBS.
15. Spin collected cells down using a centrifuge at 300g for 5 minutes.
16. Discard supernatant and resuspend cells in buffer or media.
17. Cells are ready for downstream applications.

### Validation Results

We found that cell suspensions prepared using this device vs manual dissociation have comparable cell yields and sample viability across mouse lung, kidney, and heart tissues (**Figure 6A-B**). We found that populations of immune cells like T cells and dendritic cells were not affected by a difference in isolation protocol (**Figure 6C**). Similarly, analysis of surface marker expression on these immune cell populations show similar frequencies of isolated T cells (**Figure 7A**) and comparable mean fluorescence intensity for antigen presenting marker, MHC-II in dendritic cells (**Figure 7B**) between manual and device isolation.

**Figure 6.**
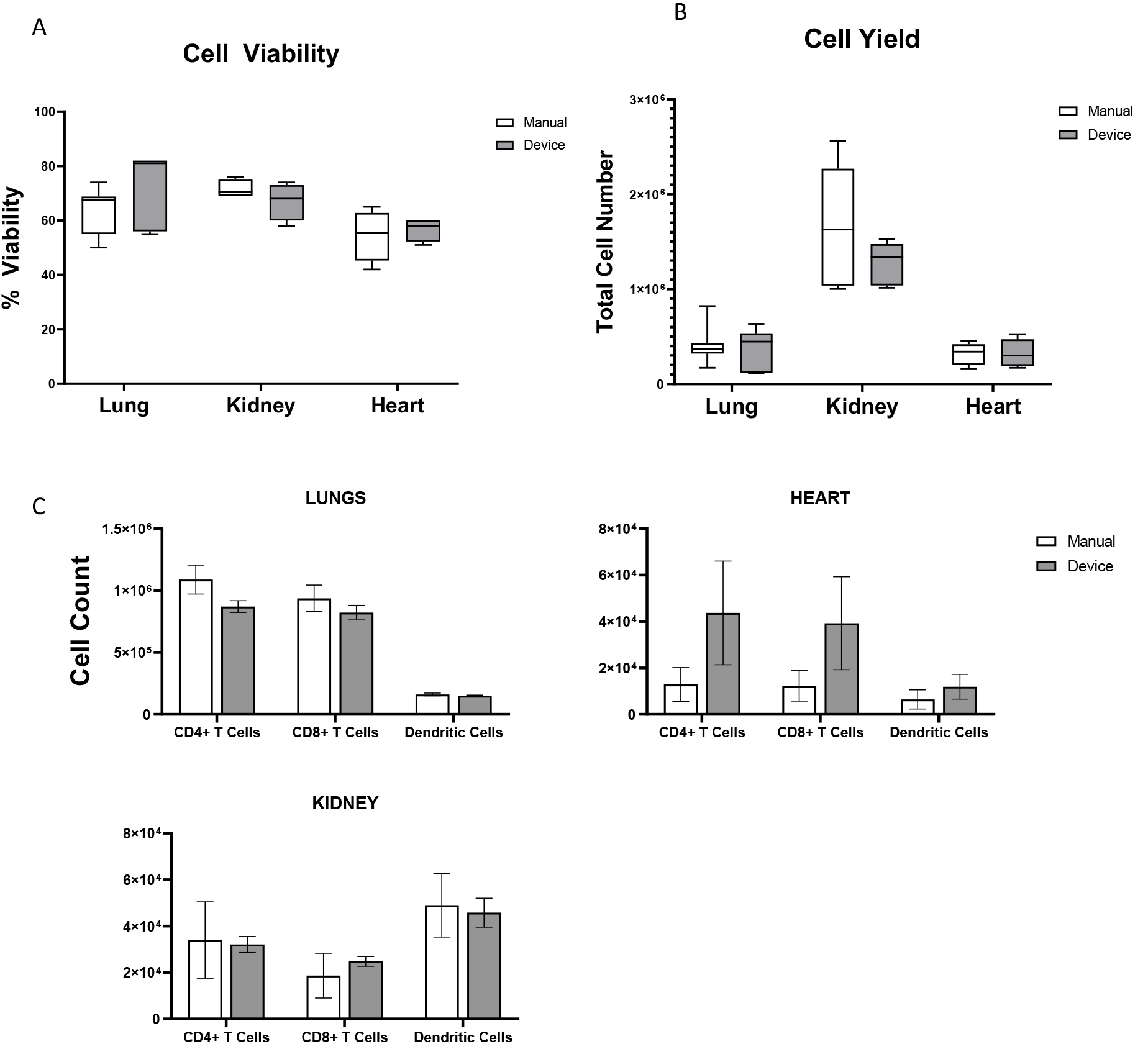
Cell Yield and Viability between Manual and Mechanical Dissociation Protocols. A) Viability and B) yeild of cells from different tissues immediately after preparation of cell suspension counted by hemocytometer before flow staining C) Cell counts of immune cell populations identified by flow cytometry. Data were analyzed using GraphPad Prism (GraphPad, La Jolla, CA). Unpaired Welch t tests were used to compare pairs of samples or groups. Differences were considered statistically significant when p < 0.05. *p<0.05, **p<0.01, ***p<0.001, ****p<0.0001.

**Figure 7.**
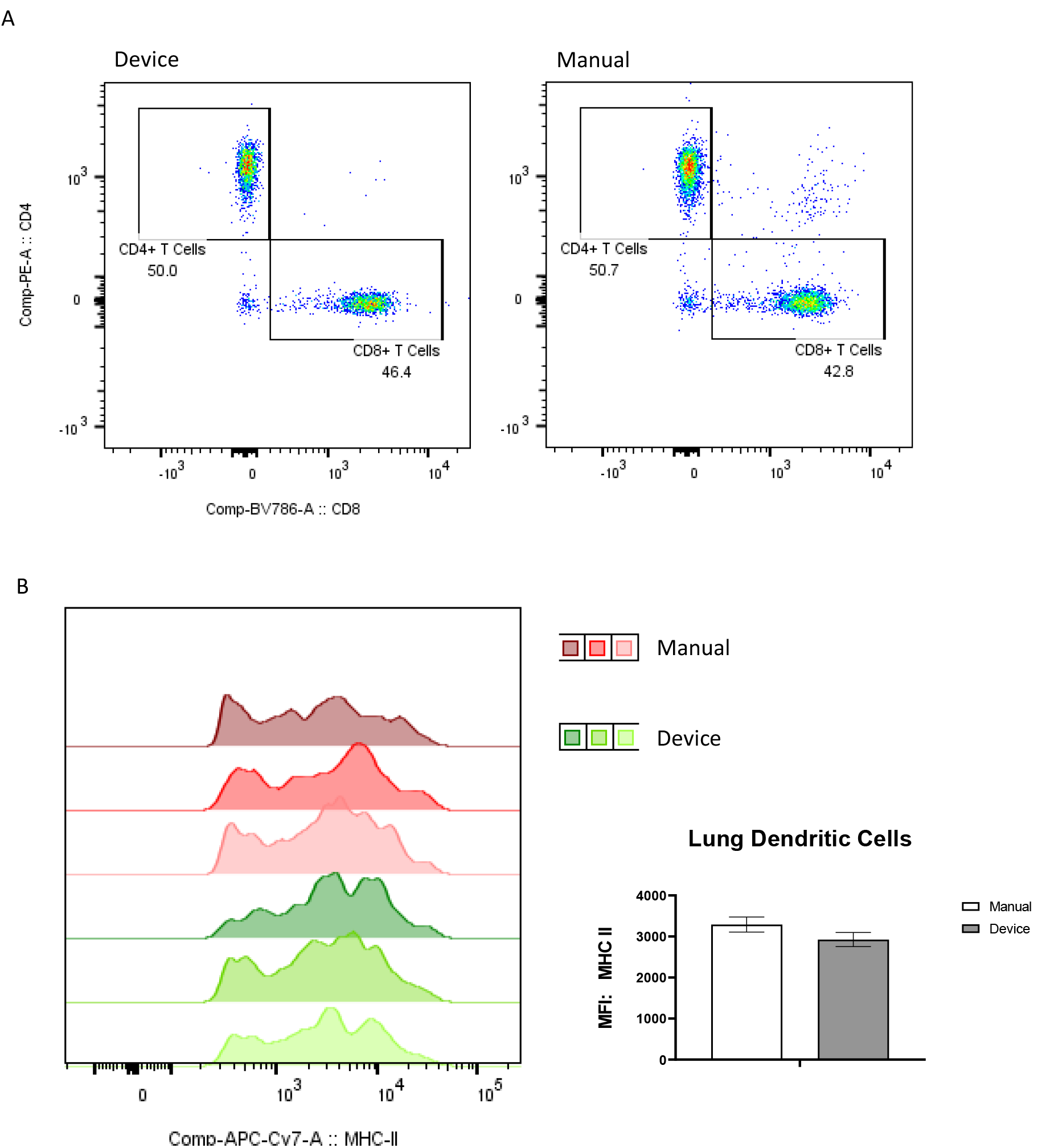
Comparison of Immunophenotypic analysis of Single Cell Suspensions between Manual vs Device Dissociation Protocols. Flow cytometry comparison of surface marker expression in single cell suspensions prepared using mechanical dissociator device vs manual dissociation. A) Lung CD4+ and CD8+ T Cell populations B) Mean fluorescence intensity (MFI) of surface MHC-II on lung dendritic cells Data were analyzed using GraphPad Prism (GraphPad, La Jolla, CA). Unpaired Welch t tests were used to compare pairs of samples or groups. Differences were considered statistically significant when p < 0.05. *p<0.05, **p<0.01, ***p<0.001, ****p<0.0001.

## CONCLUSIONS/DISCUSSION

We have designed a device that can be easily assembled in a research setting and provide single cell suspensions from whole tissues for subsequent single cell analysis. The features, though basic, are enough to meet the needs of researchers in academic settings and beyond. A key benefit of using this device is its potential to improve the preparation of single cell suspensions by reducing variability due to user inexperience. In addition, having the ability to process 12+ samples simultaneously could increase sample-to-sample consistency that may be affected by increased time required for manual dissociation protocols. In research institutions and industry settings, devices like this allow for faster and customizable tissue processing (7,8). In academic laboratories, this device could allow trainees (undergrads and graduate students) an opportunity to complete more sophisticated research by shortening the time needed to process tissues for in-vivo analyses.

### Recommendation for Non-Engineering Labs

We worked with Terrapin Works, the fabrication and prototyping center housed in the Engineering department of our university, to design and build this device. If such a resource is available at your institution, we recommend reaching out to the mechanical, electrical, and software engineers there, who should find this design relatively easy to assemble. Alternatively, we recommend reaching out to the university’s machine shop (if available) or mechanical, electrical, and/or software engineering labs local to your environment that may be able to help with the fabrication or advise on additional available resources.

### Future Improvements

We have not developed code that circumvents the manual voltage dial to instead automatically adjust the motor speed through the software. This would require hardware additions including a variable voltage regulator and more sophisticated software to incorporate this feature into the preset and custom program modes.

## STAR METHODS TEXTS

### RESOURCE AVAILABILITY

#### Lead Contact

- Further information and requests for resources and reagents should be directed to and will be fulfilled by the lead contact, Katharina Maisel

#### Materials availability

- This study did not generate new unique reagents. All materials used are widely and commercially available.

#### Data and code availability

- Viability, yield, and flow cytometry data reported in this paper will be shared by the lead contact upon request.
- All original code is available in this paper’s supplemental information.
- Any additional information required to reanalyze the data reported in this paper is available from the lead contact upon request.

## SUPPLEMENTAL ITEMS

- Cad files for building motor arrays of 2, 4, 6, 8, 10, and 12
- Software code for Arduino Nano

